# Spatiotemporal patterns of breeding challenge the successive broods model in a migratory butterfly

**DOI:** 10.64898/2026.04.01.715883

**Authors:** Aramee C. Diethelm, Cheryl B. Schultz, Stephanie R. McKnight, Emma A. Deen, Abigail M. Lehner, Emma M. Pelton, Elizabeth E. Crone

## Abstract

Migration is widely recognized as a strategy for animals to track seasonally shifting resources. Yet, seasonal and spatial dynamics of migration are challenging to study, particularly for difficult-to-track insects. Among insects, monarch butterflies (Danaus plexippus) have a well-documented fall migration, but spring breeding recolonization remains poorly understood, particularly for the western population. We conducted multi-year surveys across six regions in the western United States to characterize monarch breeding phenology and evaluate three related hypotheses: (i) the successive broods model, with discrete generations shifting activity across the breeding range, (ii) a diffusion-like expansion model with overlapping breeding periods, and (iii) a mid-summer lull model with temporary summer declines in breeding for areas near the overwintering habitat. Monarch immature presence served as an indicator of local breeding activity. Our results do not support the successive broods or mid-summer lull hypotheses.

Breeding onset occurred earlier near overwintering areas and gradually expanded north-and eastward, with sustained activity in many regions throughout the season. Termination of breeding also occurred earlier at more distant sites, resulting in longer breeding activity nearer to overwintering habitat. Immature monarch density declined with distance from overwintering areas at onset and termination, suggesting delayed colonization of peripheral regions. Together, these results support a diffusion-like expansion of breeding rather than sequential generational replacement. Western monarchs also do not initiate or terminate breeding in close synchrony with host plant availability, contrary to predictions from resource-tracking theory. These findings highlight fundamental differences between western monarch breeding dynamics and paradigms for eastern monarchs, demonstrating that a single species can employ fundamentally different spatial strategies for recolonizing its breeding range in different regions. More generally, these results distinguish insect migration from systems with direct movements between wintering and breeding habitats, and underscore the value of long-term, landscape-scale monitoring for resolving habitat use across heterogeneous environments.

## INTRODUCTION

Migration, the seasonal movement of animals to optimize survival and reproduction, is a critical strategy for many species (Alerstam et al. 2003, Dingle and Drake 2007). Migratory animals are of ecological interest due to their complex movement patterns across diverse habitats, often to track seasonally-available resources (Gatehouse and Drake 1995, Dingle 1996, Abrahms et al. 2021). Because migratory species rely on multiple distant sites throughout their life cycle, environmental changes at any single site can impact their populations across their range, increasing their vulnerability to landscape alteration (Runge et al. 2014, Abrahms et al. 2021). Understanding these vulnerabilities requires knowledge of how migratory animals use the landscape throughout the year. Advances in satellite telemetry and the Global Positioning System have made tracking large migratory vertebrates increasingly feasible (Katzner and Arlettaz 2020). Although tracking methods for invertebrates are rapidly improving (Lövei and Ferrante 2024), detailed monitoring of these smaller, less conspicuous, species remains challenging (Joo et al. 2022). Nonetheless, understanding insect migration is important because frameworks developed primarily for vertebrates may not fully apply to invertebrates, whose juveniles and adults often occupy distinct habitats (Moran 1994). Given the technological limitations of tracking systems, spatially and temporally replicated surveys offer an alternative for documenting seasonal habitat use in smaller migratory insects. In the context of widespread declines in many migratory insects (Forister et al. 2019, Wagner et al. 2021), repeated seasonal surveys may provide necessary data to better understand insect migration and the drivers of population trends in these species.

North American monarch butterflies (*Danaus plexippus* L.) are a valuable focal species for studying insect migration. Monarchs are among the most well-studied migratory insect species (Chapman et al. 2015, Chowdhury et al. 2021), with potential to inform broader questions about insect migratory behavior. Migratory monarchs produce multiple breeding generations each year that spread across the United States before late-season individuals initiate a long-distance return migration to overwintering habitat (Urquhart and Urquhart 1977, Malcolm et al. 1993). North America includes two large migratory populations: the eastern population that breeds east of the Rocky Mountains, and the western population that breeds west of the Rocky Mountains (Urquhart and Urquhart 1977, Brower 1995, Dingle et al. 2005). Decades of research have documented the fall migration of eastern monarchs to overwintering areas in central Mexico (Urquhart and Urquhart 1976, Wassenaar and Hobson 1998, Howard and Davis 2009, Flockhart et al. 2017), making this journey one of the best-understood cases of insect migration (Chapman et al. 2015). Western monarchs primarily overwinter along the California coast (Urquhart and Urquhart 1977, Tuskes and Brower 1978). Early researchers questioned whether individuals at the periphery of the western breeding range could complete the long-distance migration to coastal overwintering habitat (Brower, 1995; Malcolm, 1987; Wenner & Harris, 1993). However, further research confirmed an autumn migration from the Intermountain West and Pacific Northwest to western overwintering habitat (James & Kappen, 2021; Yang et al., 2016). Despite a detailed understanding of fall migration for North American monarch populations (e.g., Wassenaar and Hobson 1998, Yang et al. 2016, Flockhart et al. 2017, Momeni-Dehaghi et al. 2021), less is known about their spring recolonization of breeding areas, especially in the West (Pelton et al. 2019, McIntyre et al. 2024).

Compared to fall migration, spring surveys of monarchs pose greater challenges due to lower population sizes and a more diffuse geographic distribution than other seasons (Espeset et al. 2016, Erickson et al. 2023). Technology to track sparse spring populations remains limited with emerging passive monitoring methods being ineffective at such low densities (Joo et al. 2022, Lövei and Ferrante 2024). Not surprisingly, comprehensive structured spring surveys are far fewer than surveys conducted in summer (when the population is the largest; e.g., Inamine et al. 2016), fall (e.g., Culbertson et al. 2022) or winter (e.g., Vidal and Rendón-Salinas 2014, Ries et al. 2015, Schultz et al. 2017). Most knowledge of western spring recolonization patterns comes from specimen collections, citizen-science data, and inference from host plant distribution (e.g., Knight et al. 1999, Dingle et al. 2005, Dilts et al. 2019, Erickson et al. 2023, James and James 2025), while repeated seasonal field surveys designed to track recolonization over time remain uncommon (James 2016, Waterbury et al. 2019). In the East, monarch breeding begins closest to overwintering habitat (Malcolm et al. 1993). Adult eastern monarchs arrive at southern United States breeding grounds in spring, remain a few weeks, and then their offspring subsequently move farther north as local milkweed resources decline (Malcolm et al. 1993, Flockhart et al. 2013). This observed pattern aligns with the “successive broods” model, in which monarchs are proposed to progressively colonize breeding grounds across multiple generations, moving further north and east as milkweed availability shifts (Malcolm et al. 1993, Cockrell et al. 1993). This model provides a useful framework for understanding eastern monarch spring recolonization, although deviations occur, such as late-summer or fall breeding at the southern edge of the breeding range in Texas (Calvert 1999, Prysby and Oberhauser 2004, Scott et al. 2023).

Compared to eastern North America, little is known about how monarch butterflies recolonize their western summer breeding range (Pelton et al. 2019, McIntyre et al. 2024). Several factors may cause western monarch dynamics to differ from eastern monarchs. Western North America features complex topography, including high mountain ranges, which create fine-scale variation in temperature and moisture (Dilts et al. 2019). California also supports a diversity of milkweed species that span a range of ecotypes and elevations, providing host plants from early spring through late fall (Stevens and Frey 2010, Dilts et al. 2019). Year-round availability of milkweed likely enables continuous breeding for monarchs in regions like Santa Barbara County, California (Malcolm 1987, Wenner and Harris 1993), and California’s Central Valley (Yang and Cenzer 2020). Together, these factors indicate that western monarchs operate under ecological conditions distinct from those of eastern populations, warranting further investigation of their breeding phenology.

One process that often shapes seasonal movements and breeding timing in migratory insects is the tracking of spatial and temporal variation in key resources, including larval host plants and prey (Southwood 1962, Gatehouse and Drake 1995). In these systems, successive generations exploit transient pulses of suitable habitat across broad geographic gradients. For example, painted lady butterflies (*Vanessa cardui*) undertake multigenerational migrations between Africa and Europe that track seasonal increases in vegetation productivity (Stefanescu et al. 2013, 2017). Similarly, migratory hoverflies (Diptera: Syrphidae) complete long-distance seasonal movements within Europe, North America, and parts of Asia to exploit floral nectar as adults while locating ephemeral aphid outbreaks required by their predatory larvae (Reynolds et al. 2024). However, in landscapes where resources are highly spatially heterogeneous, migration may instead function as a form of bet-hedging in which insects distribute reproduction across multiple environments rather than tightly tracking a single shifting resource peak (Holland et al. 2006). Whether breeding patterns in the West are shaped primarily by milkweed phenology or by other environmental constraints remains unresolved.

Here, we investigate several biologically plausible scenarios that might predict range-wide phenology of western monarchs. In a classical successive broods model, we would expect breeding to begin and end earliest at sites closest to overwintering areas and occur progressively later at more distant sites, producing a staggered onset and termination of breeding across the range. If western monarchs follow this pattern, we would expect the first generation in California, followed by breeding further north and east, e.g., colonizing adjacent states like Nevada and Oregon in a subsequent generation, followed by Idaho, Utah, Washington and then Montana and the Canadian province of British Columbia (Table 1). In contrast to the stepwise advance implied by a successive broods model, western monarch recolonization may instead follow a diffusion-like process (Skellam 1991), in which short-distance dispersal and local reproduction would yield overlapping breeding periods across regions (Table 1). Breeding would still start earlier near overwintering areas, but instead of clear directional progression to distances far from coastal overwintering habitat, breeding activity would persist for overlapping durations. This would produce a continuous gradient of reproductive activity across the range (Table 1). As a third possibility, breeding activity could exhibit a mid-summer lull where breeding locations nearer to the overwintering habitat have two distinct periods of breeding: one early in the year and one later in the year (Table 1). One mechanism for this lull could be extreme temperatures or reduced host plant quality that temporarily limit reproduction (Yang and Cenzer 2020). This pattern is also consistent with observations that late-season breeding can occur in southern regions of central and eastern North America by eastern monarchs returning from northern breeding areas (Calvert 1999).

**Table 1:** Hypothesized breeding phenology patterns for western monarchs, showing temporal and spatial dynamics and key characteristics.

| Hypothesis | Onset of breeding | End of breeding | Notes |
| --- | --- | --- | --- |
| **Phenology in relation to calendar date and distance to overwintering sites** |  |  |  |
| Successive broods | Earlier closer to coastal overwintering sites | Earlier closer to coastal overwintering sites | Non-overlapping generations |
| Diffusion-like | Earlier closer to coastal overwintering sites | Similar or slightly later closer to overwintering sites | Breeding spreads from the coast to outer locations, but never stops near the coast |
| Mid-summer lull | Earlier closer to coastal overwintering sites | End of all breeding is later closer to overwintering sites | Two breeding periods near overwintering sites, separated by a period with no breeding (the “lull”). One breeding period at outer parts of the range. |
| **Phenology in relation to milkweed host plants** |  |  |  |
| Resource tracking | Breeding starts when milkweed is available | Breeding ends when milkweed is no longer available | Predicts similar availability of milkweed per monarch at the onset and end of breeding throughout the range. |

To evaluate these alternative hypotheses, we used multi-year monitoring to quantify monarch breeding phenology across six regions in western North America over five years.

Specifically, we asked four interrelated questions regarding the geographic patterns of monarch breeding. First, to establish the broad-scale patterns, we asked: (1) Does monarch breeding phenology differ among regions? Results from this question were used to motivate more detailed comparisons. We then evaluated predictions of the three breeding phenology hypotheses, in which we asked: (2) Does the timing of breeding onset depend on distance from overwintering habitat? Following all predicted breeding phenology models, we would expect monarch breeding to begin earlier near the overwintering habitat. (3) Does the timing of monarch breeding termination depend on distance from overwintering areas? The successive broods model predicts earlier termination near overwintering areas and later termination further away. Diffusion-like breeding would predict either similar termination times across all areas, or a slightly later end to breeding in areas near the overwintering habitat. (4) Is there a mid-summer lull in breeding near the overwintering areas? A lull is not expected under the classical successive broods model or the diffusion-like model, but would occur under the mid-summer lull model.

In addition to evaluating these spatial phenology predictions, we examined whether breeding activity is synchronized with host plant availability. We evaluated the common assumption in resource-tracking theory that monarchs should track seasonal resources during the breeding season by asking two questions: (5) Does the ratio of monarch immatures per available milkweed stem at the onset of breeding vary with distance from overwintering habitat?

Resource-tracking theory predicts that monarchs should initiate breeding in each region as host plants become available (Dingle and Drake 2007, Lemoine 2015). If monarchs are tracking resources, we expect similar immature-per-stem ratios at the onset of breeding across regions, regardless of distance from overwintering habitat. Alternatively, if monarchs are colonizing their breeding range more slowly than the rate of milkweed emergence, we expect a lower immature-per-stem ratio at the outer breeding range (i.e., more stems per monarchs). (6) Does the ratio of immatures-per-stem at the termination of the breeding season vary with distance from overwintering habitat? Resource-tracking theory predicts that monarchs should terminate breeding in each region as host plants decline or senesce. Thus, if monarchs are tracking resources, we expect similar immature-per-stem ratios at the termination of breeding across regions, regardless of distance from overwintering habitat. Conversely, a positive or negative relationship would indicate that other factors limit the termination of the breeding season.

By evaluating the three breeding phenology hypotheses (classical successive broods, diffusion-like, and mid-summer lull), we aim to clarify how western monarchs recolonize their breeding range and how the timing and overlap of breeding vary across regions. We also assess the assumption from resource-tracking theory to determine how these breeding strategies interact with regional variation in host plant availability. By resolving the seasonal breeding dynamics of a migratory insect across the American West, a region characterized by high habitat diversity and weak latitudinal gradients, this study informs our broader understanding of migration strategies.

## MATERIALS AND METHODS

### Data Collection

To understand patterns of monarch breeding across the western United States, we established study sites in six regions spanning a gradient from near overwintering habitat to more distant sites: southern California, northern California, Nevada, Idaho, Oregon, and Washington (Appendix S1: Figure S1). Surveys for monarch immatures were conducted during five years of an eight-year study period (2017–2019 and 2023–2024). Within each region, we sampled 2–5 sites (Appendix S1: Figure S1). To optimize our ability to find immature monarchs and document breeding phenology, sites were selected for suitability as monarch breeding habitat, based on host plant (milkweed) availability and variation in phenology of milkweed species to capture both early-and late-season breeding dynamics (Appendix S1: Sampling Methods; Appendix S1: Table S1). In a broad sense our goal was to sample ∼1000 stems at three locations within two sites per each region (Appendix S1: Sampling Methods). However, we added sites to make up for lack of access to some sites in some years and/or because we could not always find 1000 milkweed stems per site. Survey timing followed a latitudinal and phenological gradient to capture the full breeding season in each region, beginning with the earliest documented monarch sightings in each region: March (southern CA), April (northern CA), and May/June for distant inland and northern sites (NV, ID, OR, WA). Surveys occurred monthly until most milkweed senesced at a site (Appendix S1: Figure S2). For each survey month, sampling was conducted within a 3-week window beginning in the middle of the month, except at some sites in 2017 and 2018 where surveys were conducted up to three times per month (Appendix S1: Table S1). In 2017, additional surveys were conducted to assess the required survey effort for our hypotheses, while in 2018, surveys were added to improve detection probability in response to low counts the previous year.

During each survey, milkweed stems were identified and individually counted (details in Appendix S1: Sampling Methods). To document monarch reproductive activity at each site, we searched milkweed stems and recorded the numbers of eggs, larvae, and pupae. We focused on immature stages because they confirm reproduction within a site, whereas adult monarchs are highly mobile and their presence may not actually reflect local breeding activity. The area searched was adjusted based on site geography and milkweed patch size (more details in Appendix S1: Sampling Methods). Survey effort was kept as consistent as possible across sites and years, although survey locations shifted slightly at times due to changes in milkweed distribution or restrictions on permanent markers by landowners; the timing of surveys was also occasionally (but rarely) affected by extreme weather and logistical constraints that prevented completing surveys.

Distances between breeding and overwintering areas were calculated as great-circle distances. We compared four metrics: distance to a central overwintering site (Pismo Beach, CA; 35.138°N, 120.641°W), distance to the nearest major overwintering site, mean distance to all major overwintering sites, and mean distance to the majority of known overwintering sites in California (N = 185). Correlations among these metrics were all >0.9, showing that the relative ordering of breeding regions from closest to farthest is essentially the same regardless of which distance definition is used. Therefore, all reported distances to the overwintering habitat refer to distance to Pismo Beach, CA, in kilometers, as the central reference area used in our analyses.

## Statistical analysis

All analyses were conducted in R version 4.3.1 (R Core Team 2022).

To first evaluate the overall spatio-temporal variation in monarch breeding activity, we modeled immature monarch counts (summed across life stages) with generalized additive models (GAMs) implemented with the *mgcv* package in R (Wood 2011). Models used a negative binomial distribution with a log link, with immature monarch counts as the response. Fixed effects were survey region and a smooth term for day of year (hereafter, DOY) as well as their interaction. We included a random effect of location within sites. We also included an offset of the log-transformed number of milkweed stems surveyed to account for variation in survey effort. To assess the statistical significance of fixed effects and their interactions, we used marginal likelihood ratio tests (LRT) implemented with the *lmtest* package (Hothorn et al. 2022). This analysis did not include observations from the sites in Washington state, where breeding was observed only once during the study period (Appendix S1: Table S2), leading to problems with model convergence. This analysis informs tests of the successive broods, diffusion-like, and mid-summer lull hypotheses by characterizing temporal overlap and regional variation in breeding activity.

To estimate the onset and termination of breeding, we fit quantile regression models to immature monarch sighting dates, with the 0.1 and 0.9 quantiles representing the first and last 10% of breeding activity at each site with the *quantreg* package (Koenker 2025). We utilized quantile regression, rather than GAM-derived thresholds, to better account for the non-linear phenology of immature insects. Unlike adult abundance, which often follows a symmetric seasonal distribution, larval densities can peak abruptly upon colonization when per-capita host-plant availability is low (Yang and Cenzer 2020); therefore, the number of immatures per milkweed stem often shows a convex pattern (e.g., Pelton et al. 2019), rather than the typical concave pattern. These estimated quantiles served as spatial markers of recolonization timing and breeding cessation, providing proxies for generational dynamics by characterizing the expansion and contraction of the breeding range over time. This approach distinguishes successive-broods from diffusion-like movements by indicating whether the breeding window shifts progressively across regions or overlaps among them. Models included survey dates weighted by the number of immatures recorded on each date as the response variable and site as a categorical predictor variable. We also compared the modeled 0.1 and 0.9 quantiles to the first and last survey dates in each region to confirm that our sampling began before the onset of breeding and continued after the termination, ensuring that the temporal range of surveys captured the full breeding season.

To evaluate how onset and termination of breeding varied with distance from the overwintering region, we conducted two follow-up analyses in which the estimated onset and end dates were regressed against great-circle distance to our selected central overwintering location. The estimated onset and end dates were used as response variables, weighted by the inverse of the standard error. We chose this two-step process to account for nonindependence of replicate samples from a limited set of sites; random effects models were not feasible for these data due to the extremely low number of sightings in some regions in some years.

To evaluate whether breeding exhibited a mid-summer lull, we fitted GAMs of immature monarch counts as a function of DOY for each survey region separately. Models included a smooth term for DOY to capture seasonal phenology, a random effect for location within site and an offset for the log-transformed number of milkweed stems. We then calculated the slope and curvature of the DOY smooths at each point using the *gratia* package (Simpson 2024).

Candidate mid-summer lulls were identified as local minima, or troughs, in the GAM-predicted immature abundance, determined by points where the first derivative changed from negative to positive. In cases where the first derivative changed sign, we used the *t*-statistic of the second derivative at this point to evaluate statistical significance, and as an index of strength of convex relationship, which characterizes the curvature or sharpness of the trough in breeding activity. When there was no change in sign of the first derivative, we set the value of this index to 0.

Here, we utilized the first and second derivatives of the GAM smooths to test for significant temporal gaps in monarch breeding phenology post-regional onset. Then, following the prediction that mid-summer lulls occur primarily in regions closer to overwintering habitat, we tested for an association between the strength of the convex relationship and distance from overwintering habitat using linear regression with the strength of the convex relationship as the response variable and the distance as the predictor variable.

Finally, to assess whether western monarchs track seasonal resources, as predicted by resource-tracking theory, we used predictions from the region-specific GAMs to estimate the expected number of immatures per milkweed stem at each region’s estimated onset and termination dates. Predicted values were log-transformed and related to distance from overwintering habitat using weighted linear regression, with weights equal to the inverse of the prediction standard errors. This analysis evaluates whether monarchs synchronize their arrival and departure with host-plant availability across the western breeding range.

## RESULTS

We surveyed six regions for 3–8 months over five survey years (N = 3,056 total surveys), searching 261,315 milkweed stems and detecting 815 immature monarchs across seven milkweed species. Breeding phenology varied significantly through the season, with a strong effect of day of year (LRT: χ² = 25.1, df = 5, p < 0.001), and a significant interaction between day of year and region (LRT: χ² = 15.3, df = 3, p = 0.002; Appendix S1: Figure S3a). There was no significant effect of region alone however (Table 2). Back-transformed predicted values from the GAMs indicated generally low immature densities across all study areas with an average of < 1 immature per 300 milkweed stems (Appendix S1: Figure S4). In northern California, immature monarchs were detected in most surveys each year, whereas in Washington, immatures were detected in only a single survey across all years (Appendix S1: Table S2). In southern California, immature monarchs were detected in most surveys in 2017, 2023, and 2024, but were absent or infrequently detected in 2018 and 2019. Within each region, milkweed availability was similar across years in both the number of stems and the timing of stem availability (Appendix S1: Figure S3b, Appendix S1: Figure S5).

**Table 2:** Likelihood ratio test (LRT) results comparing generalized additive models (GAMs) for monarch butterfly (*Danaus plexippus*) immatures, testing the effects of day of year (DOY), survey region, and their interaction, with a random effect for location within sites and an offset for milkweed abundance. This analysis excludes data from Washington, where monarch immatures were observed in only one survey (see *Methods*).

| Predictor<br>Term(s) | Full Model Predictors | Reduced Model Predictors | Likelihood Ratio Test Statistics* |  |  |
| --- | --- | --- | --- | --- | --- |
| | | | $\chi^2$ | df | p-value |
| Region | DOY * Region | DOY | 25.1 | 5 | <0.001 |
| DOY | Region * DOY | Region | 1.3 | 1 | 0.3 |
| DOY ×<br>Region | DOY* Region | DOY + Region | 15.3 | 3 | 0.002 |

\* Statistics include likelihood ratio chi-square statistics ( $\chi^2$ ) and degrees of freedom (df).

Breeding onset occurred earlier near overwintering areas, with a positive relationship between breeding onset and distance from the overwintering habitat (β = 0.061 ± 0.021, t = 2.9, df = 4, p = 0.041; Figure 1a), a trend consistent with all three western monarch breeding phenology hypotheses. Termination of breeding occurred earlier in more distant regions (β = - 0.040 ± 0.014, t =-2.9, df = 4, p = 0.039), with a negative relationship between the ending of breeding and distance from overwintering habitat (Figure 1b). This is opposite the prediction of the successive broods model, which anticipates earlier termination of breeding at habitats closer to overwintering areas. Overall, the resulting breeding season was longer at sites closer to the overwintering habitat. These patterns are more consistent with diffusion-like breeding, in which overlapping breeding periods produce a continuous gradient of reproductive activity rather than discrete, staggered generations (Table 1). At all regions except southern California, our surveys spanned the entire breeding window, beginning before the estimated onset and ending after the estimated termination of breeding (Figure 1a,b). Missing the earliest and latest breeding activity at the survey sites closest to the overwintering habitat would likely make these distance associations even stronger than estimated in our models.

**Figure 1.**
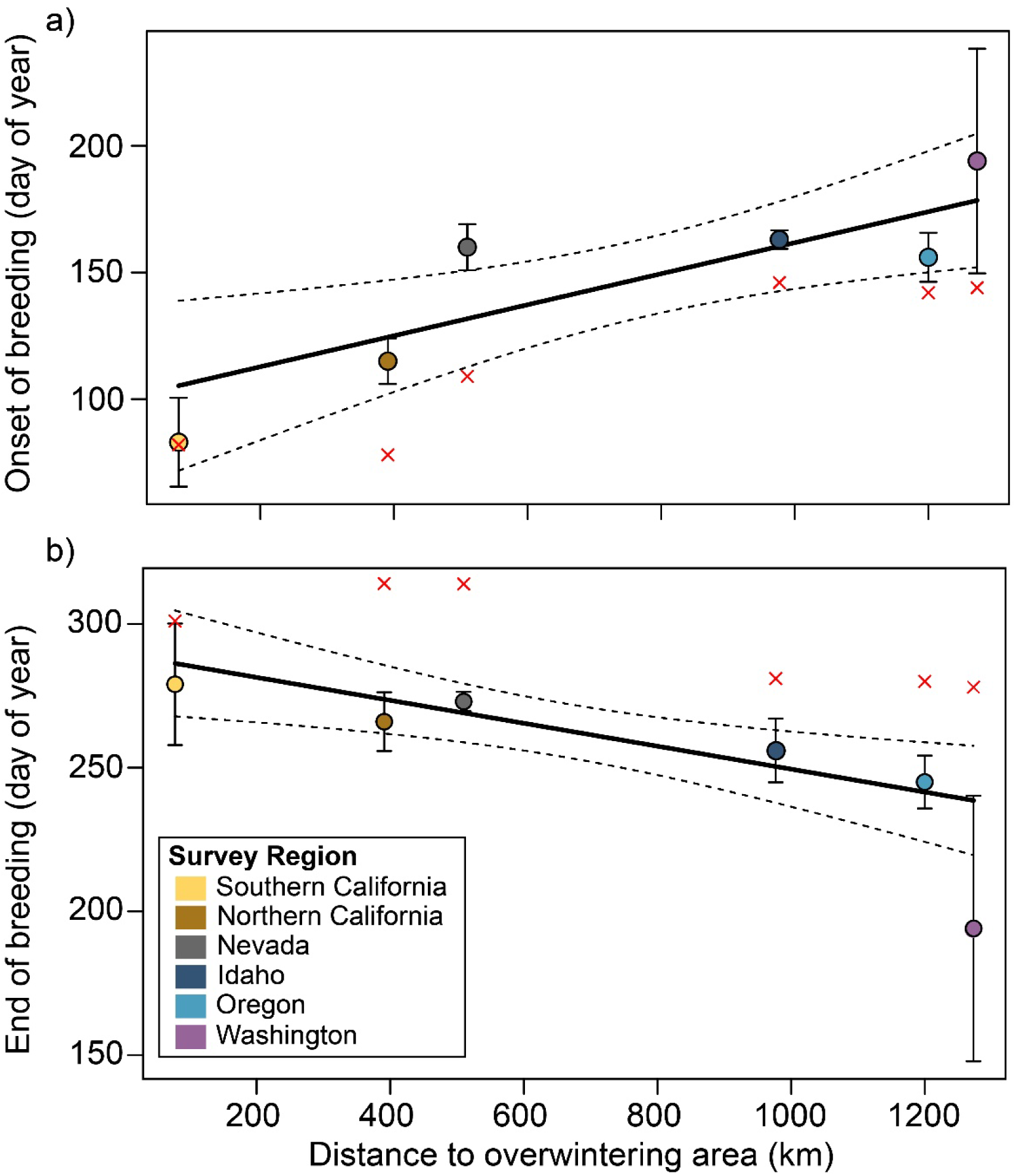
Relationships between monarch butterfly (*Danaus plexippus*) breeding phenology and distance from overwintering habitat across six regions. a) Onset of breeding (0.1 quantile of immature sighting dates). b) End of breeding (0.9 quantile). Filled circles show region-specific quantile estimates with bootstrapped standard errors. Red crosses show first (a) or last (b) survey dates for each region. Solid lines show weighted linear regressions of onset or end dates against distance to coastal overwintering habitat, and dashed lines indicate 95% confidence intervals.

We found no evidence supporting the hypothesis of a mid-summer lull in the regions closer to overwintering habitat (Figure 2). The only region exhibiting a concave mid-season pattern was Oregon, where the first derivative of the GAM crossed from negative to positive and back to negative at a statistically significant level (t = 2.05), indicating a modest mid-season decrease in breeding. Overall, mid-season dips in breeding were weak and did not follow the predicted spatial pattern of the mid-summer lull hypothesis (Figure 2). Consistent with this, support for concavity did not depend on distance to overwintering habitat (β = 0.00047 ± 0.00082, t = 0.6, df = 4, p = 0.6; Figure 2).

**Figure 2.**
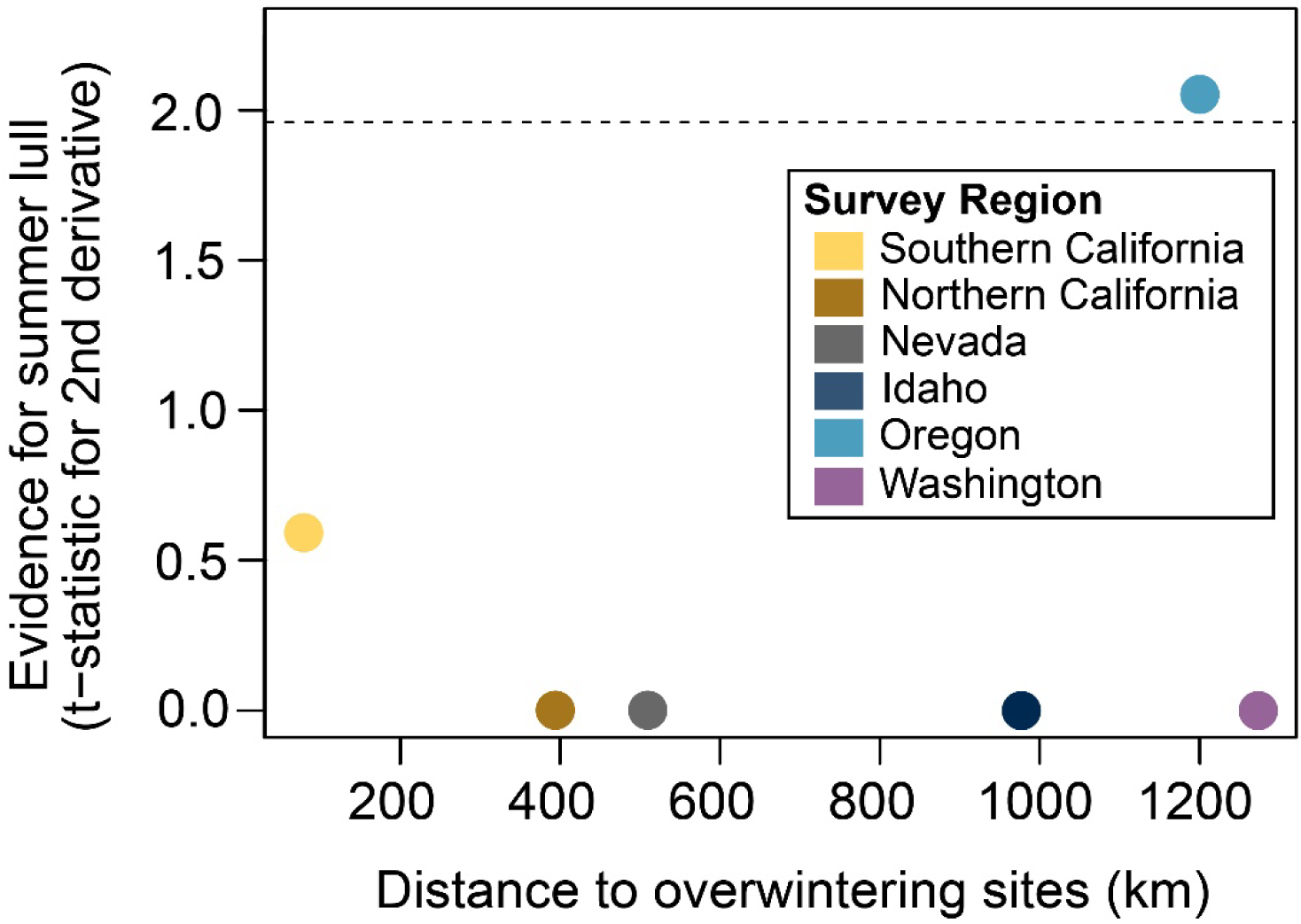
Test outcomes for a mid-summer lull in immature monarch abundance where filled circles show the t-statistic for the second derivative of the GAM-predicted abundance, with higher values indicating stronger concavity (i.e., more pronounced lull). The dashed horizontal line represents the conventional significance threshold for the t-statistic (|t| = 1.96), above which the concavity would be considered statistically significant.

Finally, we assessed milkweed availability relative to breeding onset and ending.

Immatures per stem at the onset of breeding declined with distance from overwintering habitat (β = –0.0048 ± 0.0010, t = –5.0, df = 4, p = 0.008; Figure 3a). In other words, monarch breeding at closer sites started when fewer milkweed stems were available per immature than at more distant sites. Similarly, immatures per stem at the end of the breeding period also declined with distance from overwintering habitat (β = –0.0036 ± 0.0011, t = –3.3, df = 4, p = 0.03; Figure 3b). This suggests that monarch breeding at closer sites ended when there were fewer stems per immature than at more distant sites. These patterns are inconsistent with predictions from resource-tracking theory, which expects monarchs to initiate and terminate breeding as host plant availability changes.

**Figure 3.**
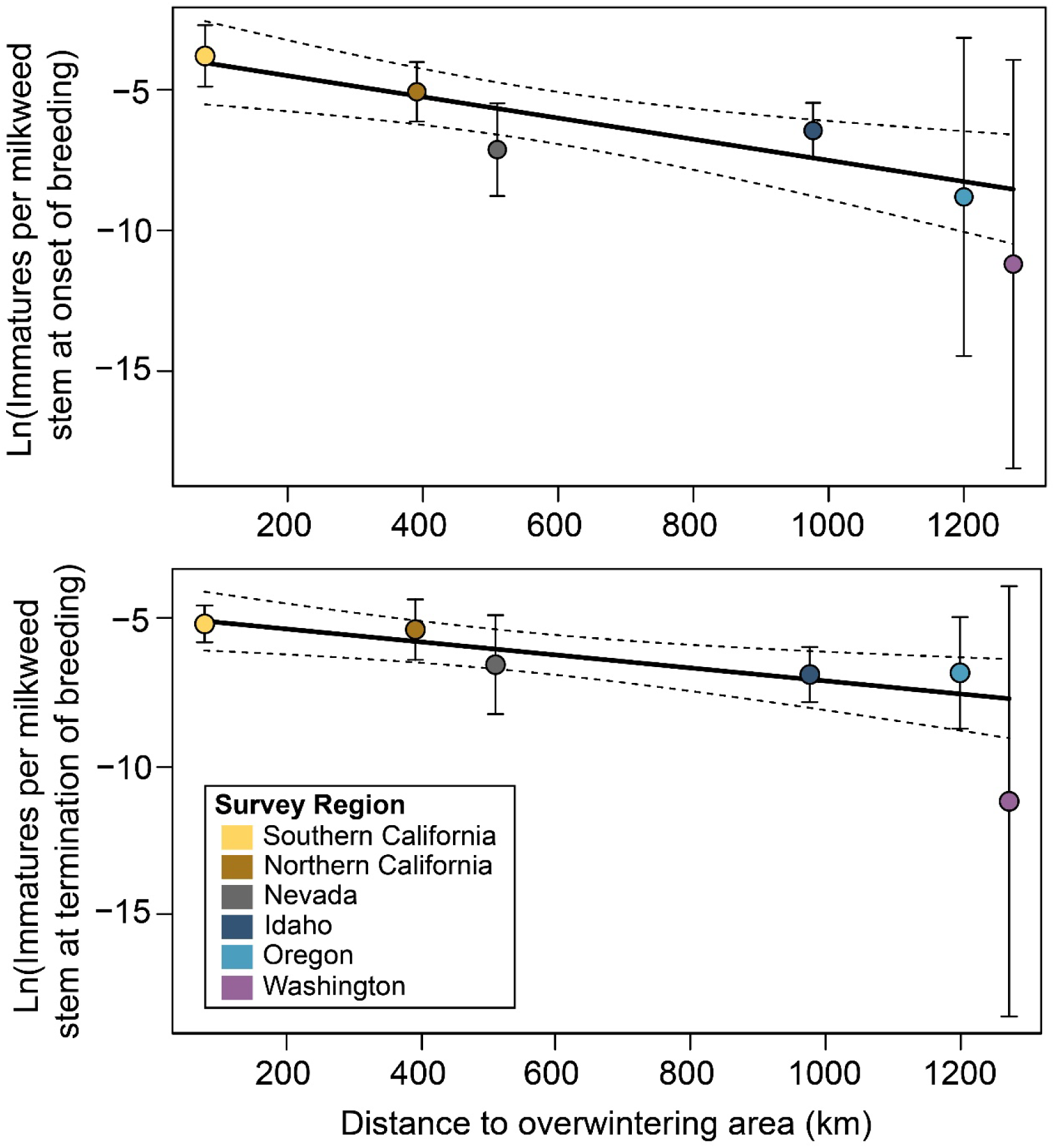
Relationships between predicted immature monarchs per milkweed stem and distance from overwintering habitat across six regions for a) onset of breeding (0.1 quantile of immature sighting dates) and b) termination of breeding (0.9 quantile), shown on a log scale. Filled circles show region-specific predictions with 95% confidence intervals. Solid lines show weighted linear regressions of predicted density against distance to coastal overwintering habitat, and dashed lines indicate 95% confidence intervals around the fitted line.

## DISCUSSION

Our findings do not support the successive broods model for western monarch breeding phenology, which predicts early-season breeding to be concentrated in areas closer to the overwintering habitat (e.g., California), ending there after the spring generation, with later generations appearing progressively farther north and east (e.g., Nevada, Oregon, Idaho), producing a staggered onset and termination of breeding across the western breeding range (Cockrell et al., 1993; Flockhart et al., 2013; Malcolm et al., 1993). Although breeding onset occurred earlier near overwintering habitat, as predicted by this model, breeding terminated earlier in more distant regions, resulting in a longer breeding season for sites closer to overwintering areas. In contrast to eastern monarch butterflies and painted lady butterflies in Europe (Malcolm et al. 1993, Stefanescu et al. 2013), western monarch butterflies do not appear to follow the generational replacement expected under a classical successive broods framework.

Instead, our results are most consistent with diffusion-like breeding model (Skellam 1991), in which breeding expands gradually across the landscape while continuing in early-season regions, producing substantial temporal overlap in breeding activity across regions. For western monarchs, breeding began earliest near the overwintering habitat in California, where monarchs remained present throughout the season, followed by a progressive north-and eastward expansion. Breeding in Nevada and Idaho began 30–60 days later than California, approximately matching the duration of an adult monarch’s life cycle or the time required for dispersal and reproduction. Classical diffusion models predict that a species’ range expansion rate depends on both movement and population growth rates (Skellam 1991, Kot 2001, Crone and Schultz 2022). Therefore, one implication of this gradual spread is that colonization of the breeding range by monarch butterflies is a process that reflects both movement and population growth. Our findings highlight that spring colonization for western monarchs differs fundamentally from other migratory species that disperse directly from wintering to breeding grounds (Dingle 1996), such as migratory songbirds (e.g., *Setophaga* spp.), Arctic caribou (*Rangifer tarandus*), and anadromous salmonids (*Oncorhynchus* spp.; Mouritsen 2018, Joly et al. 2019), as well as from monarchs’ direct fall migration back to overwintering habitat (Urquhart and Urquhart 1977, 1978).

Although our results contrast with the classical successive broods model often used to characterize breeding by eastern North American monarchs, they align with some previous research in the West. For example, Wenner and Harris (1993) documented year-round breeding near overwintering areas in Santa Barbara and proposed that monarch movements in California resulted from range expansion and contraction through random dispersal, which is consistent with our findings. However, although our results align with the general concept of an expanding and contracting population, we observed this pattern at much larger spatial scale. Additionally, our observations of summer breeding in California’s Central Valley are also consistent with observations by Yang and Cenzer (2020) in the city of Davis (∼100 km from our northern California sites). Like our findings, they documented monarch breeding throughout summer, but reported reduced mid-summer breeding success, proposing “windows of opportunity” for reproduction. We did not detect any consistent mid-summer declines, suggesting that differences between studies may reflect methodological variation, distinct pressures of monarch juveniles in urban as compared to semi-natural sites (e.g., predation, see Yang and Cenzer 2020), or interannual variation in monarch breeding phenology and juvenile success. Finally, our observed breeding phenology in Oregon suggests bivoltine reproduction (two breeding generations), and parallels recent observations by James and James (2025) across the Pacific Northwest.

Migration is often considered a strategy for tracking resource availability (Dingle 1996, Abrahms et al. 2021). Monarch distributions are commonly assumed to shift seasonally in response to host plant availability, with adults arriving when milkweed abundance is sufficient to support larval growth and departure as milkweed senesces (e.g., by late spring in the southern United States; Malcolm et al., 1993; Merlin et al., 2020; Reppert & Roode, 2018). Under resource-tracking theory, similar immature-per-stem ratios would be expected across regions at the onset of breeding, because monarchs should initiate reproduction as local milkweed becomes available. Our analyses of western monarch breeding phenology do not support this prediction.

Instead, immatures per stem at the onset of breeding declined with distance from overwintering habitat, indicating that monarchs near overwintering sites began breeding when per-capita host plant availability was low, whereas monarchs at more distant regions began breeding when milkweed was relatively more abundant per monarch. Peripheral breeding regions like northern Oregon and southern Washington were colonized well after milkweeds had fully emerged and reached substantial abundance. This observed delay in the onset of breeding relative to milkweed emergence may reflect a phenological mismatch driven by earlier milkweed emergence, later adult monarch arrival, or both, potentially mediated by climate or changes in monarch abundance. In 2017, despite lack of immatures in our survey plots in Washington, we observed fresh adult monarchs in the study area in July and August. However, this does not provide sufficient information to infer the region in which these butterflies eclosed. An additional caveat to our findings is that our Oregon monitoring sites were in the far northern part of the state and may not reflect the full range of monarch breeding phenology there. Southwestern Oregon often reports earlier and later monarch activity and may support a longer season than our sampling captured (James & James, 2025).

The observed spatial and temporal patterns of breeding likely reflect underlying dispersal dynamics. Monarch butterflies can move rapidly during fall migration (Garland & Davis, 2002; Malcolm, 1987), but colonization rates of spring breeding ranges are slower (Altizer et al. 2015, Schroeder et al. 2020), likely reflecting the balance of costs of dispersal in relation to benefits of reaching more host plants. Compared to the East, these costs could be higher in the West, where milkweed is sparse and monarch densities are low; e.g., our counts were typically one immature monarch per 300–1500 stems, in contrast to much higher counts of immature monarch butterflies in the east (one immature monarch per 1-10 stems; Stenoien et al., 2015). Furthermore, the highly heterogeneous landscape of the West may facilitate a spatial bet-hedging strategy, in which the monarch population maintains a demographic baseline in the southern breeding range while simultaneously establishing and sustaining breeding populations across the interior, thereby reducing the risk of local habitat failure (Gatehouse and Drake 1995, Holland et al. 2006). Dispersal into more distant regions may also reduce local densities of immature stages and thereby weaken density-dependent predation or parasitism (Fryxell and Sinclair 1988), potentially diluting per-capita mortality risk if natural enemies are concentrating in high-density prey patches (Jeffries and Lawton 1984).

We selected sites to represent the best available habitat to maximize detection of breeding, and therefore our sites are not random samples of the West overall in a statistical sense. Additionally, our counts do not account for losses due to predation or parasitism, so observed monarch immatures per milkweed stem may underestimate actual reproductive effort.

Nonetheless, we generally observed the highest abundance of late-season immature monarchs in both southern and northern California and Nevada, suggesting these regions may contribute disproportionately to the overwintering population during the years we surveyed. Immatures per stem at the end of the breeding period declined with distance from overwintering habitat, suggesting that monarchs farther from overwintering habitat underuse available milkweed stems relative to landscape availability. In contrast, Yang et al. (2016) found that monarchs collected at overwintering sites in 2009 most frequently originated from the northeastern inland breeding range—specifically Idaho and parts of Washington and Oregon. One possible explanation for this discrepancy is that survival to adulthood or during migration is lower in California and Nevada than Idaho, Washington and Oregon. Another explanation is that regional use of habitat may have shifted over time, potentially correlated with annual abundance at the overwintering sites. Western monarchs have both declined and become increasingly variable in abundance over past decade (Pelton et al. 2019, Xerces Society Western Monarch Count 2025), and this decline seems to be associated with a decrease in the summer breeding range size (blinded authors, *unpubl. manuscript*). A third possibility is that regions with very large milkweed stems numbers (potentially >1 million stems) such as our sites in Oregon, Idaho and Washington (James, 2016), can produce large numbers of monarchs even when immature-per-stem ratios are low. Finally, a fourth possible explanation is that monarch breeding is highly variable among years. This last interpretation echoes past work with eastern monarch butterflies: In a single-year study, Flockhart et al. (2013) initially estimated that over 50% of overwintering monarchs originated from the upper Midwest based on a single year of data, but when expanded across 38 years, contributions from that region ranged from 18–58% depending on the year (Flockhart et al. 2017).

In conclusion, we observed patterns of seasonal breeding habitat use for western monarchs that differed from established paradigms for eastern monarch butterflies. These patterns point to possible differences between eastern and western populations, and possible changes in the western population through time. Some aspects of eastern monarch breeding phenology may also be less well understood than assumed and more similar to the patterns we observed for western monarchs. From a methodological perspective, it would be interesting, though challenging, to expand our study to more sites and evaluate spatial patterns of breeding habitat use more explicitly. In some parts of the USA, such expansion could be possible through volunteer or citizen-participatory programs. In the more remote parts of the West, this is unlikely to be feasible, and a deeper understanding will require either much more extensive professional survey effort or advances in remote tracking technology. In the meantime, our results emphasize that, in spite of the fame of their fall migration, monarch butterflies do not “migrate” in a classical sense in spring. In the West, they colonize breeding habitat gradually, continuously breeding in California, and expanding into outer areas long after host plants are available. This diffusion-like pattern of recolonization may function as a form of spatial bet-hedging, distributing reproduction across heterogeneous environments and reducing both environmental risk and the potential for density-dependent mortality from natural enemy pressures. Taken together, these findings show that spring and summer recolonization can emerge from the coupling of dispersal and reproduction, producing gradual range expansion and incomplete synchrony with host resources, in contrast to classical resource-tracking or successive-broods models. Our findings also emphasize the diversity of ways in which migration can occur in insect populations.

## DATA ARCHIVING STATEMENT

All data and corresponding R code will be archived in Dryad, publicly available upon publication at: https://doi.org/10.5061/dryad.4j0zpc8q5.

## ACKNOWLEDGEMENTS

The project was motivated by the need to understand monarch breeding phenology to inform land management practices on DoD installations during the breeding season. We are especially grateful to Ryan Orndorff and Elizabeth Galli-Noble of the DoD Legacy Resource Management Program for their guidance throughout the project as well as Michael Rizo and Matt Horning with US Forest Service International Programs. We also thank the installation resource managers who facilitated access and logistics, including Ann Bedlion, Michael Bianchi, Zackary Bowers, Hodge Echeverria, Rhys Evans, Tamara Gallentine, Jessica Griffiths, Doug Grothe, Emma Hoskins, Taylor Johnson, Anna Keyzers, Claire Kurlychek, Colin Leingang, Joelle Mangelinckx, Chadwick McCready, Leslie Pena, John Philips, Kimberly Quayle, Jamieson Scott, Sydney Tomechko, Jenni Dorsey-Spitz, Gary Cottle, Donna Withers, and Lisa Weigel. Additional thanks go to collaborators at nearby reserves, refuges, and state parks—including Amy Hopperstad, Avery Hardy, Bart McDermott, Bethany Chagnon, Brenda Juarez, Bridgette Flanders, CalLee Davenport, Carl Lunderstadt, Chad Eidson, Dan Lubin, Heather Constable, Heidi Newsome, Katie Maikis, Kate McCurdy, Keely Lopez, Lamont Glass, Mary Jane West-Delgado, Nikki Evans, Sarah Trujillo, and Tanner Wolfson. Heather Constable, Chad Eidson, Amy Hopperstad, Carl Lunderstadt, and Bart McDermott—for their assistance in the field. We also acknowledge various members of the Crone and Schultz Labs for field assistance, including Michelle Boone, Lee Bennion, Sam Bussan, Cass Carroll, Emily Erickson, Samantha Hubbard, Kelsey King, Sarah Lane, Atticus Murphy, Dominick Rose, Jacob Swanson, Chelsea Thomas, and Haylie Wilcox, as well as Neal Williams for his contributions to survey work.

## CREDIT AUTHOR CONTRIBUTIONS

Aramee C. Diethelm: Methodology; Investigation; Data curation; Formal analysis; Project administration; Visualization; Writing – original draft; Writing – review & editing.

Cheryl B. Schultz: Conceptualization; Methodology; Funding acquisition; Investigation; Project administration; Supervision; Validation; Writing – original draft; Writing – review & editing.

Stephanie R. McKnight: Investigation; Data curation; Writing – review & editing.

Emma A. Deen: Investigation; Data curation; Writing – original draft; Writing – review & editing.

Abigail M. Lehner: Investigation; Visualization; Writing – original draft; Writing – review & editing.

Emma M. Pelton: Conceptualization; Methodology; Writing – review & editing.

Elizabeth E. Crone: Conceptualization; Methodology; Formal analysis; Funding acquisition; Investigation; Project administration; Supervision; Visualization; Writing – original draft; Writing – review & editing.

All authors contributed critically to manuscript development and approved the final version.

