## Appendix S1 for "Spatiotemporal patterns of breeding challenge the successive broods model in a migratory butterfly"

**Supplemental Information for:**

**Spatiotemporal patterns of breeding challenge the successive broods model in a migratory butterfly**

Aramee C. Diethelm^1*^, Cheryl B. Schultz^2^, Stephanie R. McKnight^3^, Emma A. Deen^1^, Abigail M. Lehner^4^, Emma M. Pelton^5^, and Elizabeth E. Crone^1^

^1^ Department of Evolution and Ecology, University of California, Davis, CA, 95616, USA

^2^ School of Biological Sciences, Washington State University, Vancouver, WA, USA

^3^ U.S. Fish and Wildlife Service, Port Orford, Oregon, USA^**^

^4^ Entomology and Nematology, University of California, Davis, CA, 95616, USA

^5^ The Xerces Society for Invertebrate Conservation, Portland, OR, USA

** The findings and conclusions in this article are those of the author(s) and do not necessarily represent the views of the U.S. Fish and Wildlife Service.

**Table of Contents:**

| **S1. Sampling Methods** | Page 2 |
| --- | --- |
| **Tables: Table S1** | Page 4 |
| **Tables: Table S2** | Page 7 |
| **Figures: Figure S1** | Page 8 |
| **Figures: Figure S2** | Page 9 |
| **Figures: Figure S3** | Page 10 |
| **Figures: Figure S4** | Page 11 |
| **Figures: Figure S5** | Page 13 |

**Appendix S1.** Sampling Methods

Stems were defined as shoots that were completely separate above ground, such that above-ground branches were not counted as separate stems. Stems showing more than 50% dieback were excluded from counts and they were not considered available food resources for monarch juveniles. Sampling methods included complete censuses of localized milkweed patches, linear transects, belt transects, or 2×2 m plots. In areas with fewer than 500 stems, we conducted a complete census (surveying all localized available stems). In areas with more than 500 stems, we instead employed: (1) linear transects, surveying stems within 15 cm along 150 m transect lines in areas with very dense milkweed, i.e., locations where milkweed density was effectively a carpet; (2) belt transects, covering ~250 m² between start and end points, in locations with continuous but more sparsely distributed milkweed; or (3) 2×2 m plots, in locations with patchy milkweed distribution that was not amenable to transects. All stems visibly rooted within plot were counted and checked for monarchs.

Milkweed is sparsely distributed in many parts of the West, and many of the areas in which we work have very low human population density and very little prior ecological knowledge. Therefore, locating areas with sufficient numbers of stems for annual surveys was not straightforward. Within each survey site, milkweed stems were located using a combination of historical records, conversations with researchers and managers familiar with the area and/or haphazard searches of areas we walked through while looking for known locations. For statistical analyses, we categorized broad locations within each site into areas that broadly correspond to similar habitat type, as defined by features like plant community, milkweed species present, and the presence of shade and water. For example, two sites (Stone Lakes National Wildlife Refuge in northern California and Sedgwick Reserve in southern California) included surveys in planted pollinator gardens as well as natural areas. As another example, one site (Beale Air Force Base in northern California) had three distinct locations, one of which was an open field with *Asclepias eriocarpa*, one of which was a shaded creek with *Asclepias fascicularis*, and one of which was an open creek with *A*. *fascicularis*. These were considered distinct locations within sites (Table S1). Sampling rules (described in the previous paragraph) were generally applied within each location type within each site in each year. In our associated data frame (archived in a public repository and available to reviewers during peer review), these locations column are defined as “Type_location”; note that there is a second column called “Location within site” that identified changes like the shifting of monitoring sites when necessary due to factors like mowing or inability to permanently mark sites or changes in sampling methods as milkweed populations grew or declined through time. In addition, sites and/or locations within sites were sometimes added and removed through time because, in some cases, not all locations could be surveyed in all years, due to access limitations such as fires, floods and associated erosion, and mowing of milkweed stems. In other cases, specifically Beale Air Force Base, we did not receive permission to work at the site until 2019.

**Tables**

**Table S1.** Distribution of sampling effort among sites and years.

| **Region** | **Site** | **Location within site** | **Years surveyed for immature monarchs** | **Milkweed species present** |
| --- | --- | --- | --- | --- |
| Southern California | Vandenberg Space Force Base | Coast | 2017, 2018, 2019, 2023, 2024 | *Asclepias fascicularis* |
|  | Gaviota State Park | Beach | 2017*, 2018, 2019, 2024 | *A. fascicularis* |
|  |  | Path | 2024 | *A. fascicularis* |
|  | Sedgwick Reserve | Headquarters (pollinator garden) | 2017, 2018, 2019, 2023, 2024 | *A*. *eriocarpa*, *A*. *fascicularis* |
|  |  | Reserve (natural) | 2017, 2018, 2019, 2023, 2024 | *A*. *eriocarpa*, *A*. *fascicularis* |
| Northern California | Beale Air Force Base (AFB) | A Street | 2019, 2023, 2024 | *A*. *eriocarpa* |
|  |  | Laughlin Road | 2019, 2023, 2024 | *A. fascicularis* |
|  |  | Warren Shingle | 2019, 2023, 2024 | *A. fascicularis* |
|  | Grass Valley | Field | 2024 | *A*. *eriocarpa*, *A*. *fascicularis* |
|  |  | Grass Valley | 2017, 2018, 2024 | *A*. *cordifolia*, *A*. *eriocarpa*, *A*. *fascicularis* |
|  | South Yuba River State Park | South Yuba | 2017, 2018, 2019, 2023 | *A*. *cordifolia*, *A*. *fascicularis* |
|  | Stone Lakes National Wildlife Refuge (NWR) | Pollinator garden | 2017, 2018, 2019, 2023, 2024 | *A. fascicularis*, *A. speciosa* |
|  |  | Wild area (wetland) | 2017, 2018, 2019, 2023, 2024 | *A. fascicularis*, *A. speciosa* |
| Nevada | Naval Air Station Fallon | Irrigation ditch | 2017*, 2018, 2019, 2023, 2024 | *A. fascicularis*, *A. speciosa* |
|  | Dixie Valley | Mountain | 2023, 2024 | *A*. *erosa* |
|  |  | Pond | 2017*, 2018, 2019, 2023, 2024 | *A. fascicularis*, *A. speciosa* |
|  |  | Valley | 2017*, 2018, 2019, 2023, 2024 | *A. fascicularis*, *A. speciosa* |
|  |  | Wash | 2024 | *A*. *erosa* |
|  | Stillwater NWR | Hunter | 2023, 2024 | *A. fascicularis*, *A. speciosa* |
|  |  | Stillwater | 2023, 2024 | *A. fascicularis*, *A. speciosa* |
| Idaho | Mountain Home AFB | Base | 2017, 2023, 2024 | *A. speciosa* |
|  |  | Jack’s Creek | 2019 | *A. incarnata*, *A. speciosa* |
|  | Hot Springs Rd | Hot Springs Rd | 2017, 2018, 2019, 2023, 2024 | *A. cryptoceras*, *A. speciosa* |
|  | Jack’s Creek | Jack’s Creek | 2017, 2018, 2019, 2023, 2024 | *A. incarnata*, *A. speciosa* |
|  | C.J. Strike Boat Ramp | Marina | 2017, 2018, 2019, 2023, 2024 | *A. incarnata*, *A. speciosa* |
| Oregon | Naval Weapons System Training Facility Boardman | Range | 2017, 2018, 2019, 2023, 2024 | *A. fascicularis*, *A. speciosa* |
|  | Umatilla NWR | McCormack Unit Lower | 2018*, 2019, 2023, 2024 | *A. fascicularis*, *A. speciosa* |
|  |  | McCormack Unit Mid | 2017*, 2018*, 2019, 2023, 2024 | *A. fascicularis*, *A. speciosa* |
|  |  | McCormack Unit Upper | 2017*, 2018*, 2019, 2023, 2024 | *A. fascicularis*, *A. speciosa* |
|  |  | McCormack Unit Heritage Trail | 2018*, 2019 | *A. fascicularis*, *A. speciosa* |
| Washington | Lower Crab Creek | Lower Crab Creek | 2018*, 2019, 2023, 2024 | *A. speciosa* |
|  | Yakima Training Center | Hanson | 2023, 2024 | *A. speciosa* |
|  |  | Lmuma | 2019, 2023, 2024 | *A. speciosa* |
|  |  | Sanders Meadow | 2018, 2019, 2023, 2024 | *A. speciosa* |
|  |  | Sourdough Creek | 2019 | *A. speciosa* |
| * indicates site-years with supplemental monitoring; see *Methods* in main text | | | |  |

**Table S2:** First and last day of year (DOY) when monarch (*Danaus plexippus*) immatures (eggs, larvae, or pupae) were observed at each of our survey regions in the West as pooled across all study sites for each year of the surveys. Values indicate the earliest and latest calendar days when breeding was observed based on standardized surveys (N = 3,056).

| Region | 2017 | | 2018 | | 2019 | | 2023 | | 2024 | |
| --- | --- | --- | --- | --- | --- | --- | --- | --- | --- | --- |
|  | First DOY | Last DOY | First DOY | Last DOY | First DOY | Last DOY | First DOY | Last DOY | First DOY | Last DOY |
| Southern California | 129 | 276 | - | - | 177 | 235 | 115 | 301 | 83 | 295 |
| Northern California | 119 | 277 | 115 | 296 | 114 | 296 | 118 | 250 | 108 | 274 |
| Nevada | 142 | 279 | 175 | 266 | 205 | 263 | 175 | 273 | 213 | 268 |
| Idaho | 163 | 256 | 177 | 206 | 171 | 238 | - | - | - | - |
| Oregon | 156 | 245 | 228 | 253 | - | - | - | - | 235 | 235 |
| Washington | - | - | - | - | - | - | 194 | 194 | - | - |

**Figures**

**Figure S1**: Map of focal study regions (N = 6) across the western United States, with the dotted line representing the Continental Divide, which separates eastern and western monarch butterfly (*Danaus plexippus*) breeding zones along the Rocky Mountains. Circles represent milkweed patches within sites where surveys were conducted (e.g., transects, 2 × 2 m plots, etc.), colored by the number of years surveyed during the study period. Some survey locations may have shifted slightly between years due to changes in milkweed availability, disturbances, or site access. Insets show detailed views of study areas within each region.

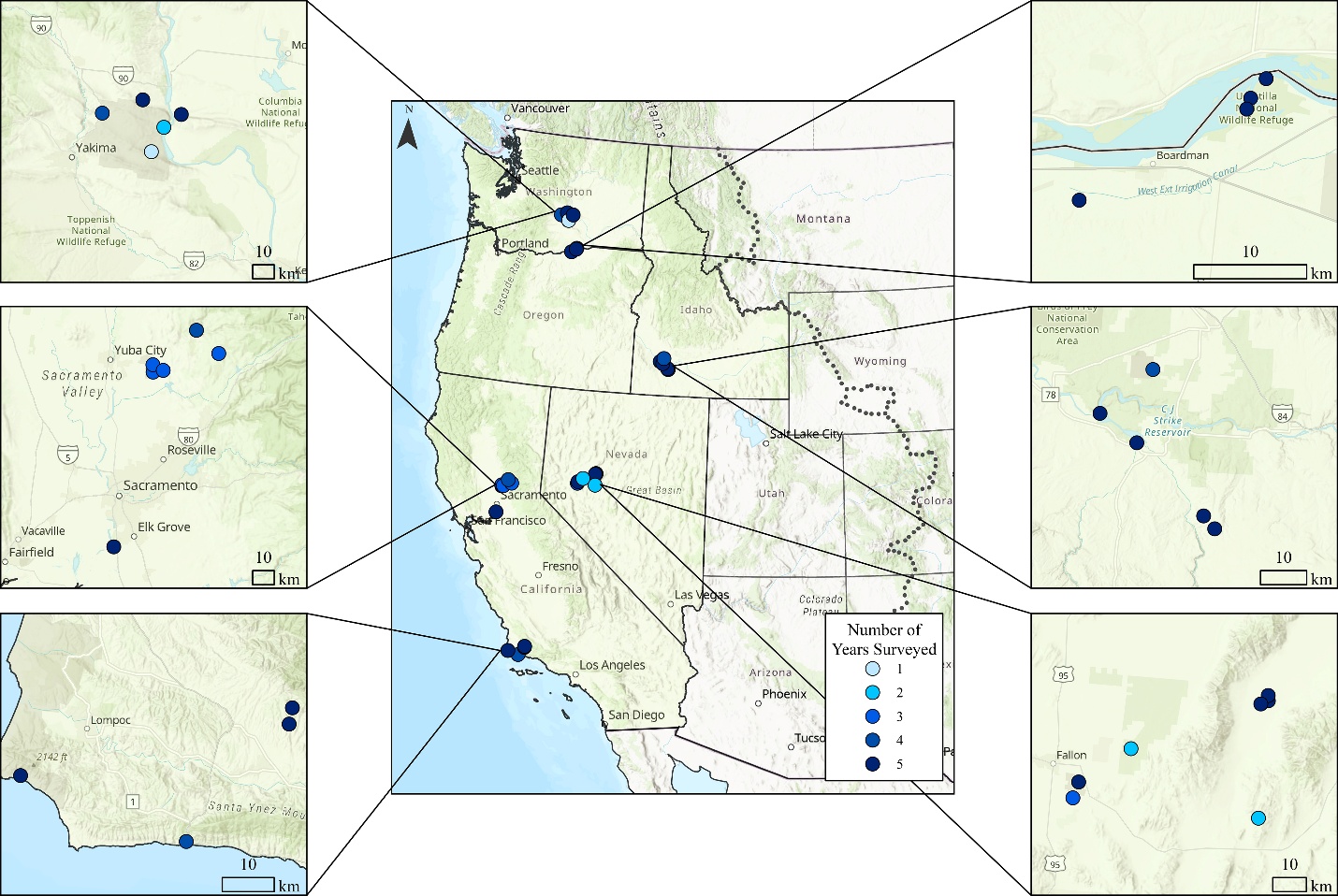

**Figure S2.** Timing of monarch butterfly (*Danaus plexippus*) breeding surveys (N = 3,057) by year and region (ordered by proximity to overwintering grounds), indicating months with (green) and without (grey) observed breeding activity.

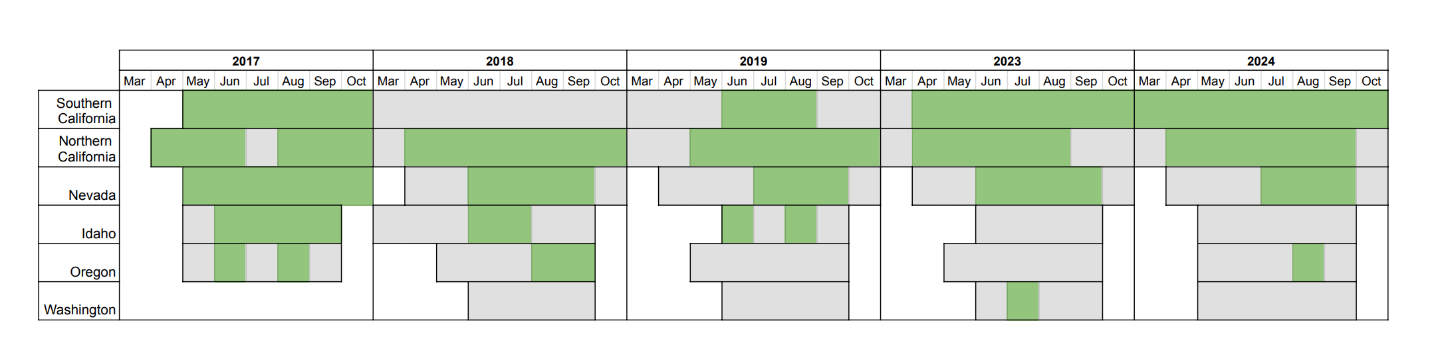

**Figure S3.** (a) Monarch breeding and (b) milkweed phenology across regions and years. In both panels, regions are ordered from closest to furthest from overwintering areas. Horizontal lines are the 10^th^, 50^th^, and 90^th^ percentiles of abundance through time, and width is proportional to the abundance at each point in time. In (a) vertical dashed lines represent times when surveys were conducted but no monarch breeding was observed.

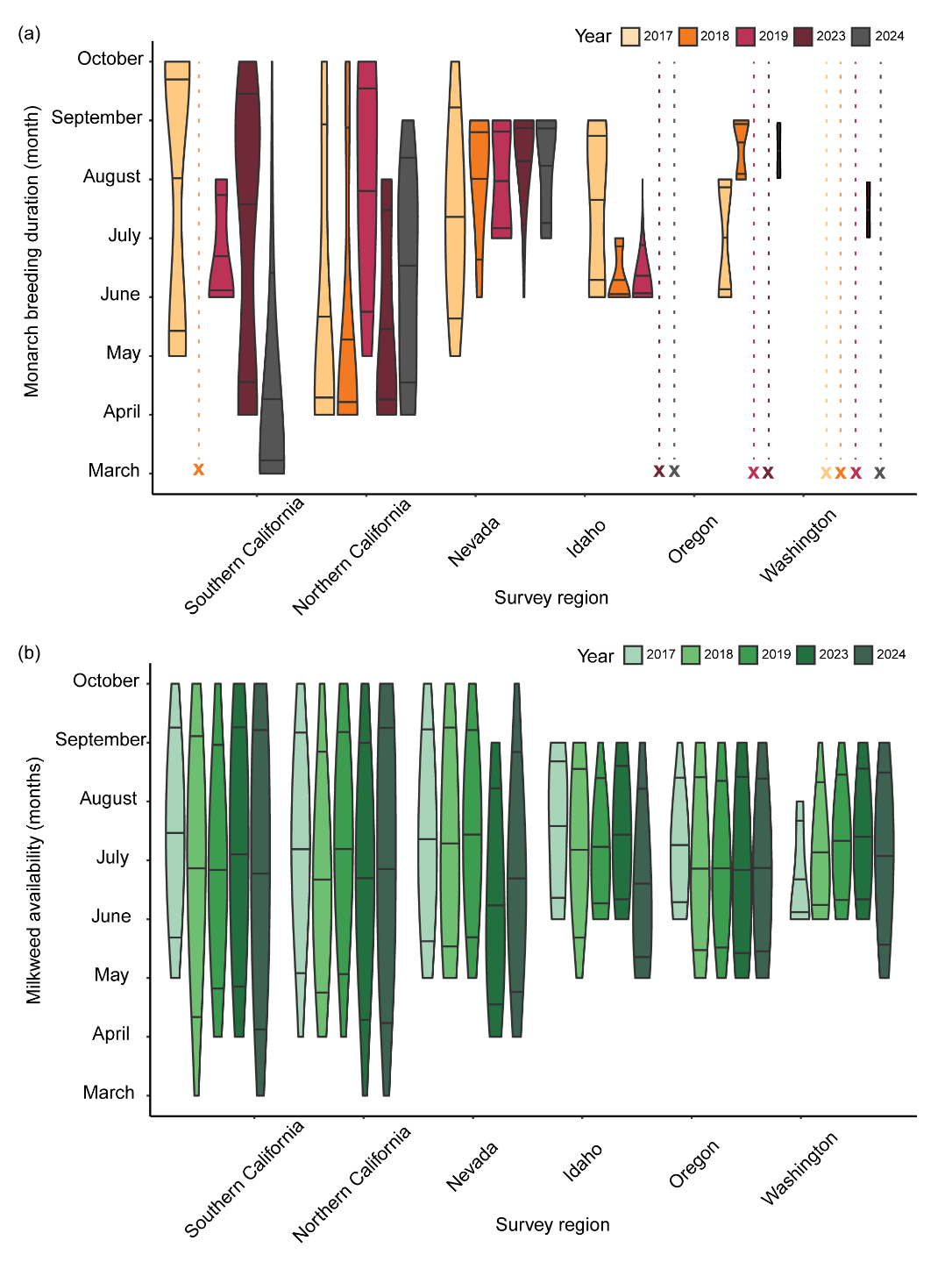

**Figure S4.** Data from surveys with observations pooled across all years (2017–2019 and 2023–2024) per survey region: southern California, northern California, Nevada, Idaho, Oregon, and Washington. For each region (each row), the x-axis is scaled to fit the data range for the three displayed metrics: 1) a ratio of monarch (*Danaus plexippus*) juveniles (eggs, larvae, and pupae) observed per milkweed (*Asclepias*) stems searched (circles), 2) raw milkweed stem abundance (triangle), and 3) raw juvenile monarch abundance (diamonds). Solid lines represent back-transformed predictions from generalized additive models (GAMs) with smooth terms for day of year (DOY) and a random effect for site type (habitat). Gray dashed lines indicate 95% confidence intervals. The x-axes are scaled to fit all regions based on the data range per graph-type.

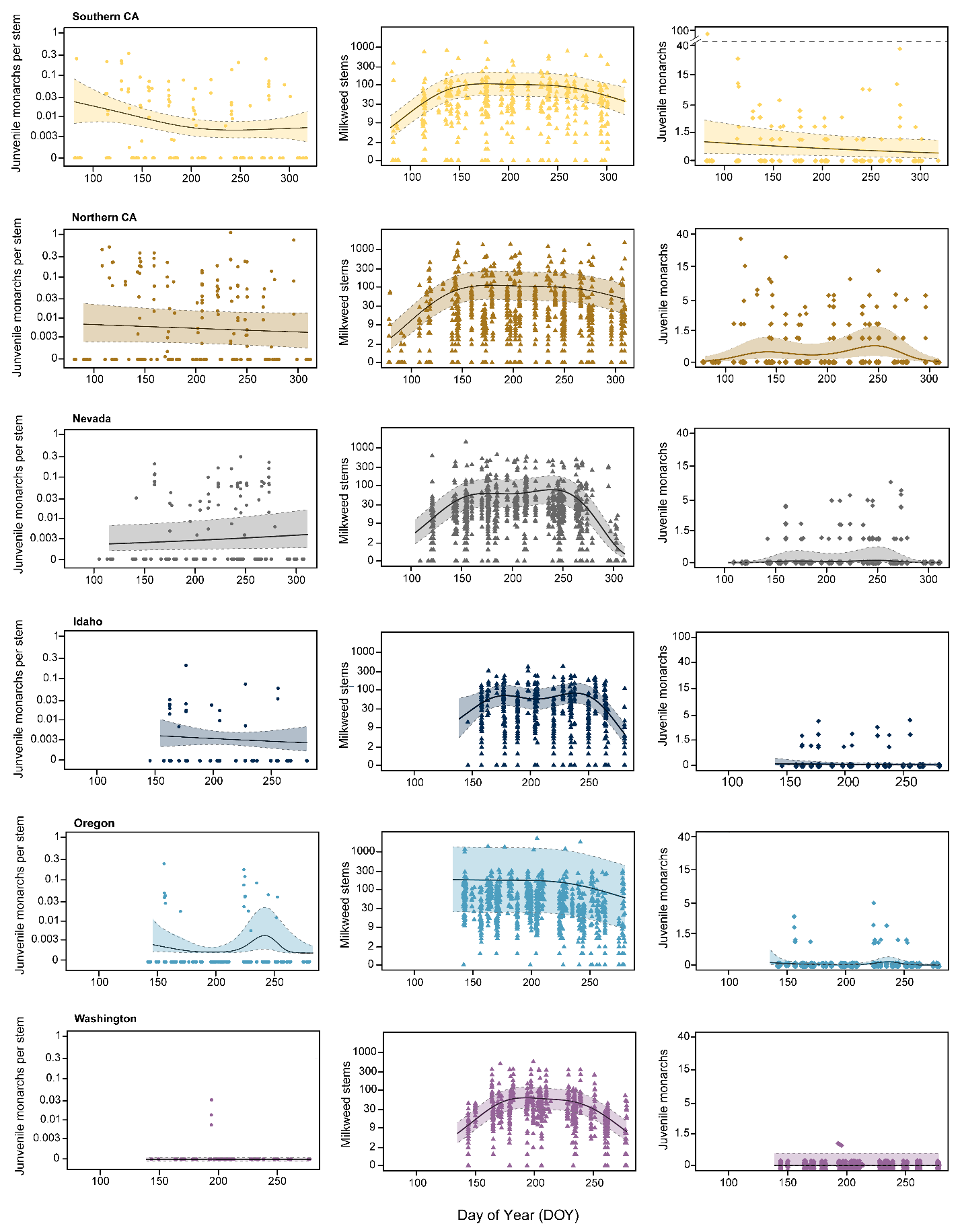

**Figure S5:** Availability and abundance of milkweed species across regions and years, ordered from nearest to farthest from overwintering sites, where the violin plot widths represent relative stem abundance and horizontal lines indicate the 10th, 50th, and 90th percentiles. This graph includes only months in which we conducted surveys, and so does not represent the full phenology of milkweed, especially in outer states like Oregon and Washington.

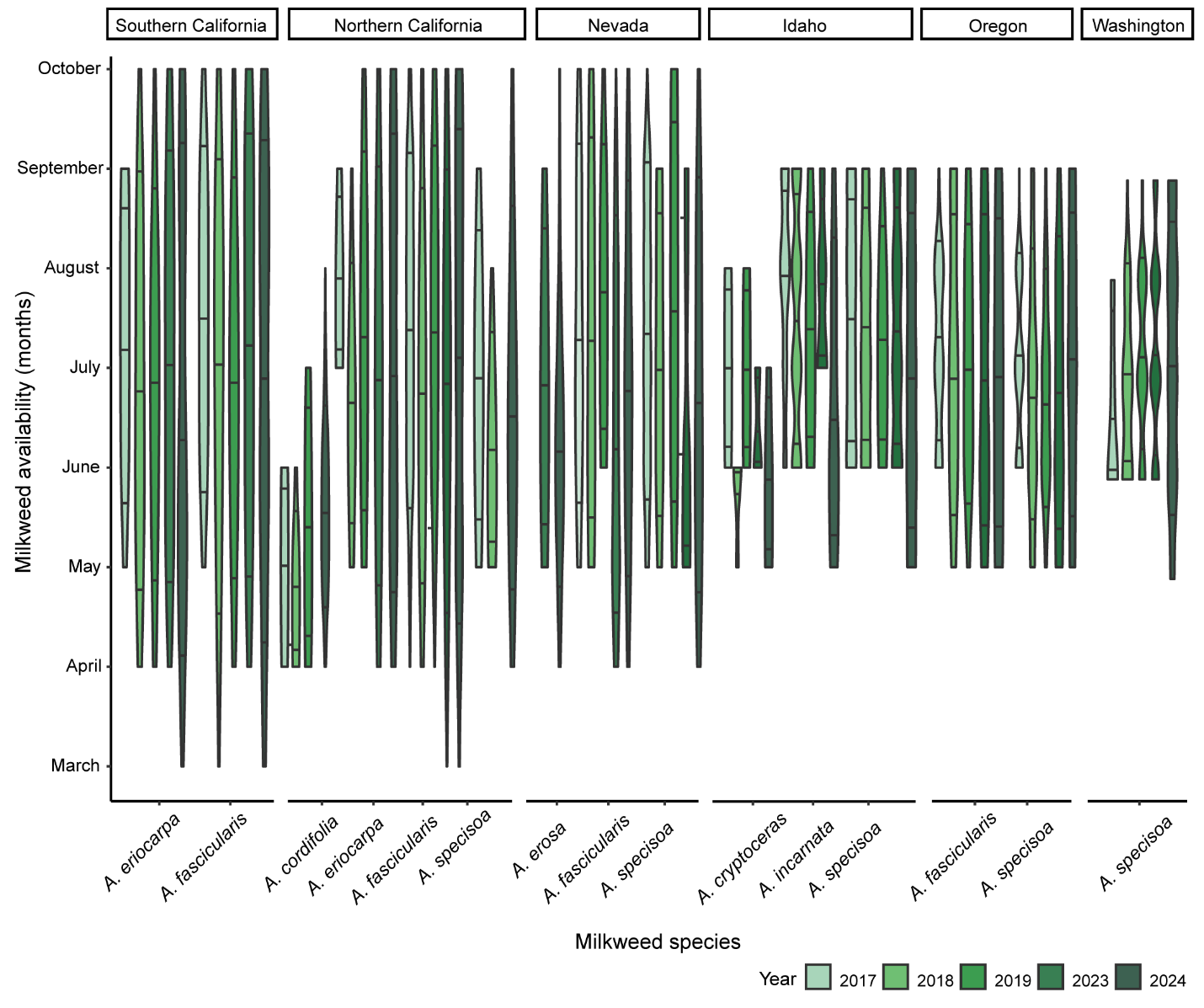
